# Global Data Fitting Enables Kinetic Dissection of Cullin-RING Ligase-Catalyzed Ubiquitylation: From Substrate Priming to Poly-ubiquitin Chain Formation

**DOI:** 10.64898/2026.11.04.764329

**Authors:** Nicholas Purser, Jerry Li, Daniel C Scott, Brenda A Schulman, Tyler L Dangerfield, Gary Kleiger

**Affiliations:** Department of Chemistry and Biochemistry, University of Nevada, Las Vegas, Las Vegas, NV, USA; Department of Structural Biology, St. Jude Children’s Research Hospital, Memphis, TN 38105, USA; Department of Molecular Machines and Signaling, Max Planck Institute of Biochemistry, Martinsried, Germany

## Abstract

The discovery and characterization of the ubiquitin system was founded on classical enzymology approaches that established fundamental principles of protein ubiquitylation. Cullin-RING ligases (CRLs) represent the largest class of human ubiquitin ligases, yet detailed enzymology of these multi-subunit assemblies has lagged behind other systems such as nucleic acid polymerases. Here, we employed pre-steady state kinetics and global data fitting using KinTek Explorer to systematically investigate the kinetic parameters controlling CRL-dependent ubiquitin transfer. Using a well-characterized *in vitro* reconstituted system exploring CUL2-dependent CRLs, we modeled substrate priming, showing that the system is highly amenable to global fitting of kinetic data. The results show that product dissociation from the CRL is significantly slower than substrate dissociation, explaining previously reported slow steady-state ubiquitylation rates. Extending this framework to poly-ubiquitin chain formation revealed catalytic efficiencies 15–42-fold greater than substrate priming. Comparative analysis of CUL1-dependent and CUL2-dependent systems showed remarkably similar kinetic parameters, including the *K*_*m*_ of ubiquitin-carrying enzyme for CRLs, despite differences in substrate receptor and enzyme architecture. We further examined drug-induced neo-substrate ubiquitylation using the pomalidomide-CRBN-SALL4 system, revealing atypical kinetics characterized by rapid ubiquitin transfer paired with an elevated *K*_*m*_, ultimately yielding catalytic efficiencies comparable to natural substrates. The findings establish pre-steady state kinetics and global data fitting as powerful enzymological tools for dissecting CRL-dependent ubiquitylation across different CRL families and substrate contexts, with important implications for understanding both physiological ubiquitylation and therapeutic targeted protein degradation strategies.

## Introduction

The post-translational modification ubiquitylation is controlled by a three-enzyme system that catalyzes the sequential transfer of ubiquitin to target proteins (1,2). The first enzyme, E1 (ubiquitin-activating enzyme), uses ATP to adenylate ubiquitin and then transfers it to a cysteine residue in the E1 active site (3,4). The second enzyme, E2 (ubiquitin-conjugating enzyme), accepts ubiquitin from E1 through a transthioesterification reaction, forming an E2∼ubiquitin thioester intermediate (5-9) (the thioester linkage is denoted throughout by the symbol ∼).

E3 ubiquitin ligases supply the specificity of the system. They recognize protein substrates bearing degrons, short sequence motifs that are frequently exposed by a post-translational modification such as phosphorylation (10). Many E3 ligases function by bringing substrates into proximity with E2∼ubiquitin (11), catalyzing transfer of ubiquitin from the E2 thioester, typically onto a Lys residue of the substrate.

Attachment of the first ubiquitin to a substrate Lys, termed substrate priming, initiates conjugation. The primed ubiquitin then serves as an acceptor for further additions. The C-terminus of each incoming ubiquitin is joined through an isopeptide bond to a lysine of the preceding ubiquitin, building a poly-ubiquitin chain. This iterative chain extension is often carried out by E2–E3 machinery not involved in priming and frequently with distinct kinetics, a difference that is central to the present work.

Ubiquitin presents seven internal Lys residues (K6, K11, K27, K29, K33, K48, and K63) together with its N-terminus, each a potential attachment point for the next ubiquitin (12). Different E2–E3 pairs build chains linked preferentially

Global Data Fitting Enables Kinetic Dissection of Cullin-RING Ligase-Catalyzed Ubiquitylation through particular lysines, producing topologically distinct poly-ubiquitin chains, and this encodes biological function. For example, K48-linked chains, the first type identified, target proteins to the proteasome and provided the original link between ubiquitin modification and regulated protein degradation (13). Because the rate of ubiquitin transfer appears to differ among acceptor lysines during both priming and chain formation, the specificity of ubiquitylation is to a large degree a kinetic phenomenon. This premise motivated the quantitative approach to Cullin-RING ligase (CRL)–dependent ubiquitylation described here.

CRLs are the largest family of human E3 ubiquitin ligases (14,15) and are estimated to mediate roughly 20% of ubiquitin-dependent protein degradation in cells (16). Each is a multi-subunit assembly built on a cullin scaffold (CUL1– CUL7 and CUL9 in humans) and organized around three functional modules: the cullin, which nucleates the complex; RBX1 or RBX2, a RING-domain subunit that recruits and helps position the E2∼ubiquitin for catalysis (17,18), and a substrate-receptor module that selects the target. Distinct cullins pair with distinct receptor families so that a single cullin–RBX catalytic core can be directed at thousands of substrates: CUL1 with F-box proteins (>70 in humans) through SKP1 adaptors (19), CUL2 with VHL- and SOCS-box proteins (20), CUL4 with DCAFs (21), and CUL5 with SOCS-box proteins. Covalent attachment of the ubiquitin-like protein NEDD8 to the cullin serves as an activation switch, sharply increasing the rate of catalysis (17,22-24) and enabling tight regulation of CRL activity (25-29).

The catalytic output of a CRL also depends on which ubiquitin-carrying enzyme it recruits. At least six E2 enzymes act with human CRLs, including UBE2D-family members specialized for substrate priming (30), UBE2R-family members that preferentially build K48-linked chains and prime certain CUL2-dependent substrates (31,32), and UBE2G1, which also builds K48-linked chains (33). In addition, the RING-between-RING (RBR) E3 ligases ARIH1 and ARIH2 act as priming enzymes for several CRLs (18,34). Because these two are themselves E3 ligases rather than canonical E2s, we use the inclusive term ubiquitin-carrying enzyme (UCE) for all eight enzymes that deliver ubiquitin through CRLs (6). The availability of multiple UCE–CRL combinations, each suited to priming or to chain formation, seems to enable a single architecture to ubiquitylate thousands of proteins efficiently.

Mechanistic enzymology has a long and productive history in the ubiquitin field (35-42). Dissection of the E1 reaction began in the 1980s (4) and was followed by detailed characterization of E2 and E3 enzymes, including multi-subunit E3 ligases (2). Enzymological studies of CRLs themselves, beginning in the early 2000s, established several principles that frame the present work: that substrate priming is frequently rate-limiting (43), that additional UCEs operate with CRLs (33), that NEDD8 conjugation upregulates activity (17,22,44), and that specific features of UCEs and E3s govern the specificity of ubiquitin transfer (17,31,45).

A landmark advance came in 2009, when Pierce and Deshaies applied rapid-quench, pre-steady-state kinetics to CRL-dependent ubiquitylation, providing the first quantitative framework for substrate priming and for the individual transfer steps of poly-ubiquitin chain formation (46). That work shaped more than a decade of our own efforts to dissect CRL mechanism by transient kinetics. Progress toward more complex mechanisms has nonetheless been limited, less by any single curve-fitting method than by a reliance on simplified kinetic models, which typically account only for substrate and product dissociation beyond ubiquitin transfer, and on conventional error analysis that can report deceptively tight uncertainties even where individual rate constants are poorly constrained by the data (52).

Here we combined global fitting of CRL-dependent ubiquitylation in KinTek Explorer (47-49) with confidence-contour analysis to extract maximal mechanistic information from single-turnover and steady-state time courses, resulting in a substantially more rigorous model of CRL-dependent ubiquitylation. This model quantifies substrate affinity in the full enzymatic context, precisely estimates catalytic efficiencies across diverse UCE-CRL combinations, reveals mechanistic principles conserved throughout the CRL family, and demonstrates that the catalytic efficiency of molecular-glue-induced neo-substrate ubiquitylation matches that of natural substrates despite vastly different *k*_*cat*_ and *K*_*m*_ values.

## Results

### Substrate Priming Kinetics in a CRL2 System

We began by reconstituting substrate priming with a minimal, fully defined CRL2 system in which every component was present at a known concentration. The ligase comprised the substrate receptor VHL bound to the Global Data Fitting Enables Kinetic Dissection of Cullin-RING Ligase-Catalyzed Ubiquitylation adapter proteins ELONGIN B and ELONGIN C, the cullin CUL2 modified with the activator NEDD8, and RBX1 (^N8^CRL2^VHL^), paired with the ubiquitin-carrying enzyme UBE2R2, a validated physiological partner of neddylated CRL2 (31). The substrate was a HIF1α-derived peptide bearing a single physiological ubiquitylation site (50). To enforce mono-ubiquitylation and simplify the kinetic analysis, a ubiquitin variant lacking the major chain-forming lysine (K48R ubiquitin) was used together with a substrate containing only one acceptor lysine (Figure 1a and Supplementary Figure S1a). Reactions were initiated in a rapid quench-flow instrument by mixing a pre-formed complex of radiolabeled substrate (S0) and ligase (CRL) with UBE2R2. A 100-fold excess of unlabeled substrate was introduced upon initiation of the reaction to prevent rebinding of dissociated radiolabeled substrate, enabling observation of a single enzymatic turnover (Figure 1b).

**Figure 1.**
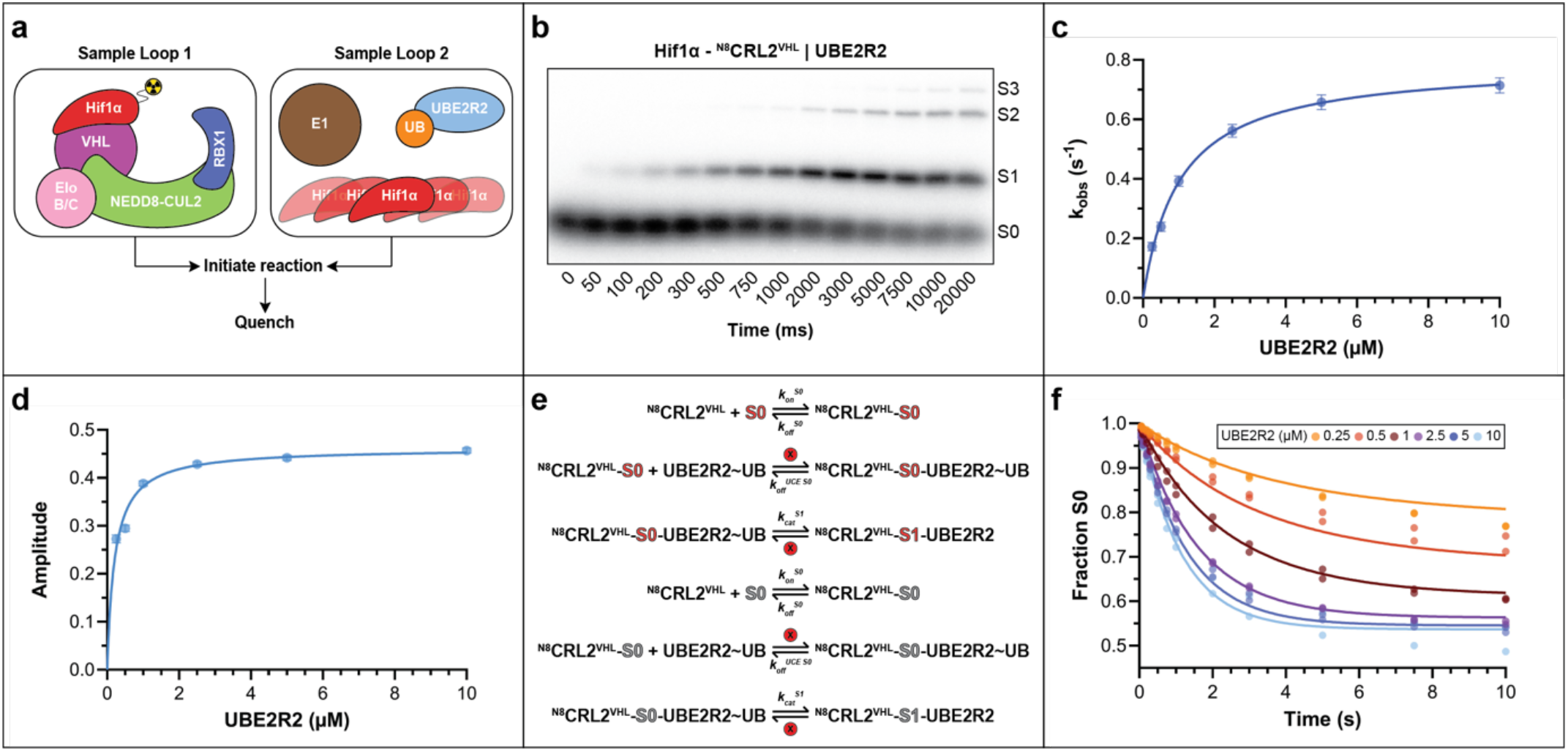
A novel approach to modeling CRL2-dependent substrate priming by ubiquitin-carrying enzyme UBE2R2 using global data fitting. (a) Reaction scheme for pre-steady state ubiquitylation. Neddylated CRL2^VHL^ with radiolabeled Hif1α peptide (S0) was mixed by quench-flow with UBE2R2 thioesterified to K48R ubiquitin under single-encounter conditions. (b) Autoradiogram showing primary conversion of S0 substrate to S1 product at 10 μM UBE2R2. (c) Graph showing the exponential observed rate constant (*k*_*obs*_) for S0 disappearance as a function of UBE2R2 concentration. (d) Graph showing the amplitude of S0 depletion at equilibrium as a function of UBE2R2 concentration. (e) Kinetic model showing rate constants for radiolabeled S0 (red) and UCE binding to neddylated CRL2^VHL^ and ubiquitin transfer (*k*_*cat*_,^*S*^,s) to S0, forming S1. Circled ‘x’ indicates rate constants with locked values. Unlabeled excess S0 (grey) promotes single-encounter conditions between labeled S0 and CRL. (f) S0 depletion as a function of time and UBE2R2 concentration, with best-fit lines from KinTek Explorer based on the model (e). Duplicate data points are shown from n=2 technical replicates. Abbreviations: UB, ubiquitin; EloB/C, ELONGIN B/C; ^N8^CRL2^VHL^, neddylated CUL2-RBX1 in complex with VHL-EloB/C substrate receptor; UCE, ubiquitin-carrying-enzyme.

We initially examined the single-turnover results and the observed rate of product formation and its amplitude (defined as the fraction of product formation at equilibrium) by nonlinear curve fitting. Analysis of the rate and amplitude informs whether the experiment can constrain rate constants such as those governing substrate and UCE binding to the CRL as well as the chemistry of ubiquitin transfer. Every trace corresponding to titration of UBE2R2 (Supplementary Figure S1b-f) was well described by a single exponential, and the observed rate increased hyperbolically with UBE2R2 concentration, saturating at high enzyme concentration (Figure 1c). This behavior is the kinetic signature of a two-step mechanism in which UBE2R2 binds in a rapid equilibrium that precedes a slower, rate-limiting chemical step and is supported by previous kinetic measurements (51). Thus, the hyperbola defines the *K*_*m*_ of UCE for the CRL and the maximal rate of ubiquitin transfer, *k*_*cat*_.

Yet another observation that guided the kinetic model was that the amplitude of traces corresponding to increasing UBE2R2 concentration rose hyperbolically rather than reaching a common endpoint (Figure 1d). We note that any reaction with an irreversible chemical step should display convergence of product formation on the same endpoint given sufficient time regardless of UBE2R2 concentration. As such, here the observed amplitude dependence instead reports an equilibrium established before the chemical step, the partitioning of S0 between free and CRL-bound states, that is progressively captured as higher UBE2R2 concentrations accelerate the forward reaction.

Guided by these features, the full family of traces were fit by numerical integration of the rate equations in KinTek Explorer (47,48) to a model comprising reversible substrate binding (CRL + S0 ⇌CRL·S0), a rapid-equilibrium UBE2R2-binding step, and an irreversible ubiquitin-transfer step (Figure 1e). Because the association rate for a rapid equilibrium binding event cannot be resolved, the value for *k*_*on*_ was fixed for UBE2R2 at the diffusion limit (100 µM^−1^ s^−1^), such that the fit reported the ratio of its dissociation to association rates (*K*_*m*_) rather than either rate individually. Global fitting described the data well (Figure 1f), and confidence-contour (FitSpace) analysis confirmed that each floated parameter was well constrained by the data (Supplementary Figure S1g-j; an excellent review of confidence-contour analysis can be found at (52)). As expected, the amplitude dependence constrained substrate binding, defining a pre-steady-state dissociation constant for S0 of 0.5 µM for ^N8^CRL2^VHL^, whereas the hyperbolic rate dependence defined a Michaelis constant of *K*_*m*_ = 1.7 µM for UBE2R2 and a catalytic rate *k*_*cat*_^S1^ = 1.0 s^−1^, resulting in a catalytic efficiency *k*_*cat*_^S1^/*K*_*m*_ of 0.6 µM^−1^ s^−1^ (Tables 1 and 2).

**Table 1.** Values for kinetic rate constants of Hif1α, Cyclin E, and SALL4(378-435) substrates and their respective CRL partners.

| **CRL** | **Substrate** | **UCE** | **Parameter** | **Best Fit** | **95% CI** |
|---|---|---|---|---|---|
| <sup>N8</sup>CRL2<sup>VHL</sup> | Hif1α | UBE2R2 | *k<sub>on</sub><sup>S0</sup>* | 0.24 µM<sup>-1</sup>s<sup>-1</sup> | 0.14–0.38 |
|  |  |  | *k<sub>off</sub><sup>S0</sup>* | 0.12 s<sup>-1</sup> | 0.068–0.17 |
|  |  |  | *k<sub>on</sub><sup>UCE S0</sup>* | 100 µM<sup>-1</sup>s<sup>-1</sup> | Locked |
|  |  |  | *k<sub>off</sub><sup>UCE S0</sup>* | 170 s<sup>-1</sup> | 134–224 |
|  |  |  | *k<sub>cat</sub><sup>S1</sup>* | 1.0 s<sup>-1</sup> | 0.89–1.2 |
|  |  |  | *k<sub>cat</sub><sup>S0</sup>* | 0 s<sup>-1</sup> | Locked |
|  |  |  | *k<sub>on</sub><sup>UCE S1</sup>* | 100 µM<sup>-1</sup>s<sup>-1</sup> | Locked |
|  |  |  | *k<sub>off</sub><sup>S1</sup>* | 0.020 s<sup>-1</sup> | 0.0082–0.039 |
| **CRL** | **Substrate** | **UCE** | **Parameter** | **Best Fit** | **95% CI** |
| <sup>N8</sup>CRL1<sup>FBXW7</sup> | Cyclin E | ARIH1 | *k<sub>on</sub><sup>S0</sup>* | 0.21 µM<sup>-1</sup>s<sup>-1</sup> | 0.14–0.32 |
|  |  |  | *k<sub>off</sub><sup>S0</sup>* | 0.060 s<sup>-1</sup> | 0.040–0.087 |
|  |  |  | *k<sub>on</sub><sup>UCE S0</sup>* | 100 µM<sup>-1</sup>s<sup>-1</sup> | Locked |
|  |  |  | *k<sub>off</sub><sup>UCE S0</sup>* | 166 s<sup>-1</sup> | 122–266 |
|  |  |  | *k<sub>cat</sub><sup>S1</sup>* | 2.6 s<sup>-1</sup> | 2.1–3.6 |
|  |  |  | *k<sub>cat</sub><sup>S0</sup>* | 0 s<sup>-1</sup> | Locked |
|  |  |  | *k<sub>on</sub><sup>UCE S1</sup>* | 100 µM<sup>-1</sup>s<sup>-1</sup> | Locked |
|  |  |  | *k<sub>off</sub><sup>S1</sup>* | 0.010 s<sup>-1</sup> | <0.029 |
| **CRL** | **Substrate** | **UCE** | **Parameter** | **Best Fit** | **95% CI** |
| <sup>N8</sup>CRL4<sup>CRBN</sup> | SALL4 | UBE2D3 | *k<sub>on</sub><sup>S0</sup>* | 0.63 µM<sup>-1</sup>s<sup>-1</sup> | 0.47–0.82 |
|  |  |  | *k<sub>off</sub><sup>S0</sup>* | 0.15 s<sup>-1</sup> | 0.12–0.19 |
|  |  |  | *k<sub>on</sub><sup>UCE S0</sup>* | 100 µM<sup>-1</sup>s<sup>-1</sup> | Locked |
|  |  |  | *k<sub>off</sub><sup>UCE S0</sup>* | 3900 s<sup>-1</sup> | >3100 |
|  |  |  | *k<sub>cat</sub><sup>S1</sup>* | 46 s<sup>-1</sup> | >37 |
|  |  |  | *k<sub>cat</sub><sup>S0</sup>* | 0 s<sup>-1</sup> | Locked |

*k<sub>on</sub><sup>S0</sup>* represents the rate constant for association of substrate and CRL; *k<sub>off</sub><sup>S0</sup>* the rate of substrate dissociation from CRL. *k<sub>on</sub><sup>UCE SX</sup>* represents the rate of UCE association with CRL, and ‘x’ reflects the number of ubiquitins attached to substrate; *k<sub>off</sub><sup>UCE SX</sup>* represents the rate of UCE dissociation from CRL. *k<sub>cat</sub><sup>S1</sup>* and *k<sub>cat</sub><sup>S0</sup>* represent the forward and reverse rate constants for ubiquitin transfer from UCE to substrate. *k<sub>off</sub><sup>S1</sup>* is the rate constant for dissociation of primed substrate from the CRL.

**Table 2.** *K*_*d,app*_ for substrate and CRL, pre-steady state *K*_*m*_ of UCE for CRL during substrate priming, and substrate priming catalytic efficiency.

| CRL | Substrate | UCE | Parameter | Best Fit | 95% CI |
|---|---|---|---|---|---|
| <sup>N8</sup>CRL2<sup>VHL</sup> | Hif1α | UBE2R2 | *K<sub>d,app</sub>* | 0.48 µM | 0.44–0.51 |
|  |  |  | *K<sub>m</sub>* | 1.7 µM | 1.3–2.3 |
|  |  |  | *k<sub>cat</sub><sup>S1</sup>*/*K<sub>m</sub>* | 0.58 µM<sup>-1</sup>s<sup>-1</sup> | 0.50–0.73 |
| <sup>N8</sup>CRL1<sup>FBXW7</sup> | Cyclin E | ARIH1 | *K<sub>d,app</sub>* | 0.29 µM | 0.26–0.32 |
|  |  |  | *K<sub>m</sub>* | 1.7 µM | 1.3–2.7 |
|  |  |  | *k<sub>cat</sub><sup>S1</sup>*/*K<sub>m</sub>* | 1.6 µM<sup>-1</sup>s<sup>-1</sup> | 1.3–1.9 |
| <sup>N8</sup>CRL4<sup>CRBN</sup> | SALL4 | UBE2D3 | *K<sub>d,app</sub>* | 0.24 µM | 0.23–0.26 |
|  |  |  | *K<sub>m</sub>* | 39 µM | >32 |
|  |  |  | *k<sub>cat</sub><sup>S1</sup>*/*K<sub>m</sub>* | 1.2 µM<sup>-1</sup>s<sup>-1</sup> | 1.1–1.3 |

Dissociation constants of substrate for CRL (*K<sub>d,app</sub>*), pre-steady-state Michaelis constants of UCEs for CRLs (*K<sub>m</sub>*), and catalytic efficiencies (*k<sub>cat</sub>*/*K<sub>m</sub>*) were estimated from the corresponding best-fit elementary rate constants.

The fitted *k*_*cat*_^S1^ of 1.0 s^−1^ greatly exceeds the steady-state ubiquitylation rates reported for this system (44,53,54), which necessitates that a step after chemistry limit multiple-turnover catalysis. It was reasoned that slow release of the ubiquitylated product from ^N8^CRL2^VHL^ could impose this limit and was tested directly by measuring steady-state ubiquitylation in the presence and absence of unlabeled competitor substrate (Figure 2). Fitting these steady-state data globally with the single-turnover experiment (Supplementary Figure S2a-f) required extending the model to include binding of the ubiquitylated product, with association and dissociation rate constants *k*_*on*_^S1^ and *k*_*off*_^S1^. Lacking a direct measurement of the product association rate, *k*_*off*_^S1^ was linked to *k*_*off*_^S0^ while the value for *k*_*off*_^S1^ was allowed to float. Confidence-contour analysis showed that the dissociation rate was well constrained by the combined dataset (Supplementary Figure S2g). Product release (*k*_*on*_^S1^) was approximately 4-fold slower than release of unmodified substrate (*k*_*on*_^S0^; Table 1), demonstrating that the single ubiquitin slows product dissociation sufficiently to make it the predominant rate-limiting step under steady-state conditions.

**Figure 2.**
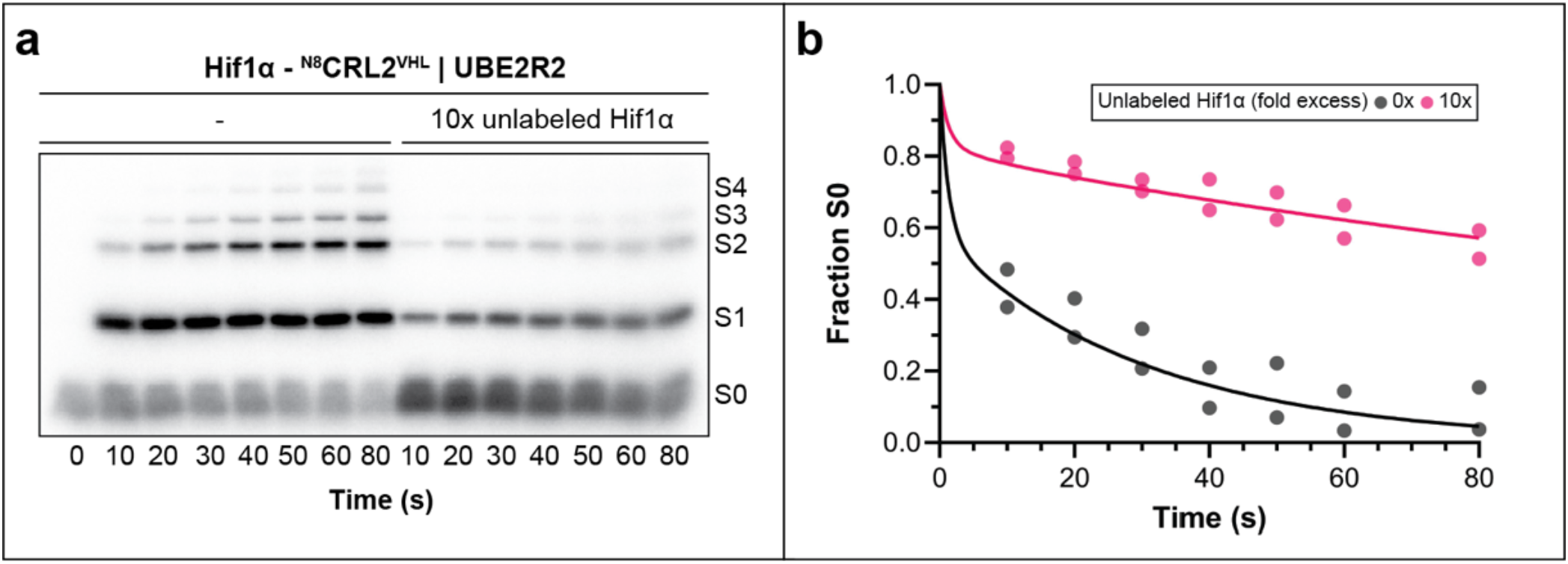
Steady-state ubiquitylation enables estimation of product dissociation kinetics, indicating product inhibition of ^N8^CRL2^VHL^ by Hif1α-UB. (a) Representative autoradiogram for single-turnover (left half) and multi-turnover (right half) ubiquitylation time courses. (b) Substrate (S0) depletion kinetics for single-turnover (gray) and multi-turnover (magenta) reactions over time and the fit to the model (see Supplementary Figure S2a). Duplicate data points are shown from n=2 technical replicates.

### Poly-ubiquitin Chain Formation Proceeds with Dramatically Elevated Catalytic Efficiency

Having defined the kinetics of priming, the successive transfers that build a poly-ubiquitin chain were addressed next. We followed the same single-turnover strategy but used wild-type ubiquitin and a mono-ubiquitylated Hif1α peptide (S1) as the starting substrate (Supplementary Figure S3a), so that each transfer lengthened the chain by one ubiquitin. SDS-PAGE resolved the growing products, and the fractional abundance of each species (S1 through S5) was quantified over time.

The priming model was expanded to incorporate processive ubiquitin transfer that begins with S1 and generates S2 through S5, incorporating substrate and product binding for each species, UBE2R2∼ubiquitin binding and dissociation, ubiquitin transfer to each intermediate, and rebinding of apo-UBE2R2 to support successive rounds of catalysis (Figure 3a). To keep the number of floated parameters commensurate with the information in the data, all reverse ubiquitin-transfer rates were fixed to zero, the UBE2R2-related association steps were constrained to the diffusion limit (100 μM^-1^ s^-1^) while allowing for the rate of UBE2R2∼ubiquitin dissociation from the CRL to float, and substrate/product binding rate constants were linked across chain lengths (see methods). Pre-steady-state time courses collected across a range of UBE2R2 concentrations provided a good fit to this model, and all floated parameters were well constrained by confidence-contour analysis (Figure 3b–e and Supplementary Figures S3b–i and S4).

**Figure 3.**
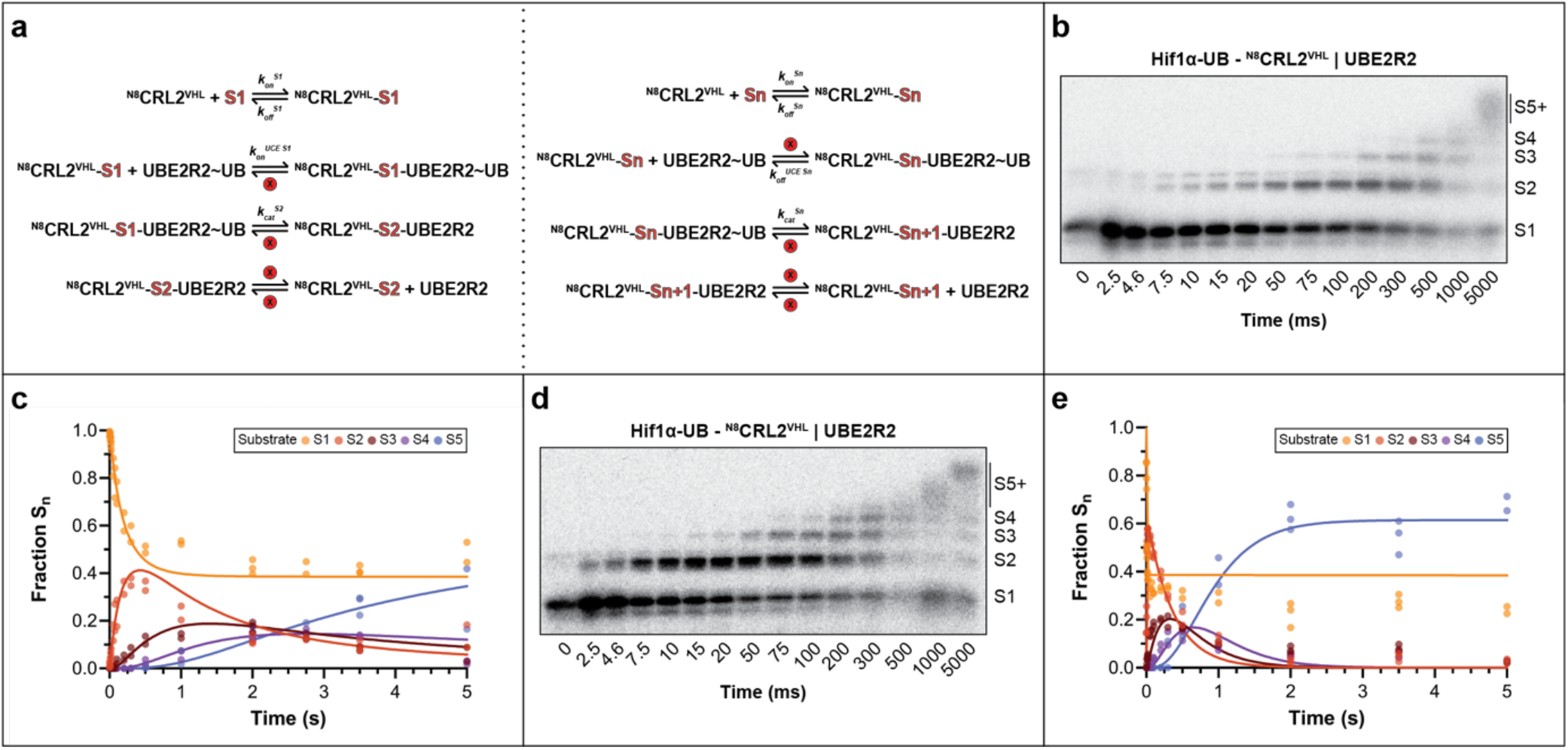
Global data fitting enables catalytic efficiency estimation for poly-ubiquitin chain formation steps. (a) Kinetic model highlighting rate constants for iterative poly-ubiquitin chain formation. The scheme on the left shows molecular events during elongation of the poly-ubiquitin chain from mono-ubiquitylated substrate S1 to product S2. The one on the right shows the general case where n varies from 2 to 4 here. Circled ‘x’ indicates rate constants with locked values. (b) Representative autoradiogram at 0.25 μM UBE2R2 showing S1 conversion to poly-ubiquitylated products. All non-overlapping product bands were quantified. (c) S1 loss and appearance of products S2–S5 with best-fit lines from KinTek Explorer. (d,e) Same experiments with 10 μM UBE2R2: autoradiogram (d) and corresponding kinetic traces with best-fit lines (e). Duplicate data points are shown from n=2 technical replicates.

The catalytic efficiency (*k*_*cat*_/*K*_*m*_) for transfer to the primed substrate S1 was 23 µM^−1^ s^−1^, and the subsequent transfers (S2→S3, S3→S4, and S4→S5) proceeded with efficiencies between 8.1 to 13 µM^−1^ s^−1^ (Table 3). The Michaelis constants governing UBE2R2 during chain formation were uniformly low (0.27–0.36 µM), approximately 6-fold tighter than during priming (1.7 µM). As a result, chain-formation efficiencies exceeded the priming efficiency by approximately 15-to 42-fold, confirming the notion that the UBE2R2–CRL pair is kinetically specialized to elongate chains (30,39,43,46).

**Table 3.** Pre-steady state *K*_*m*_, *k*_*cat*_, and catalytic efficiencies for poly-ubiquitin chain formation.

| CRL | Substrate | UCE | Parameter | Best Fit | 95% CI |
|---|---|---|---|---|---|
| <sup>N8</sup>CRL2<sup>VHL</sup> | Hif1α-UB | UBE2R2 | *k<sub>+1</sub>* | 0.0041 | 0.0038–0.0044 |
|  |  |  | *k<sub>cat</sub><sup>S2</sup>/K<sub>m</sub><sup>S2</sup>* | 23 µM<sup>-1</sup>s<sup>-1</sup> | 20–28 |
|  |  |  | *K<sub>m</sub><sup>S3</sup>* | 0.36 µM | 0.17–0.70 |
|  |  |  | *k<sub>cat</sub><sup>S3</sup>* | 2.9 s<sup>-1</sup> | 2.2–3.8 |
|  |  |  | *k<sub>cat</sub><sup>S3</sup>/K<sub>m</sub><sup>S3</sup>* | 8.1 µM<sup>-1</sup>s<sup>-1</sup> | 5.6–13 |
|  |  |  | *K<sub>m</sub><sup>S4</sup>* | 0.33 µM | 0.085–0.87 |
|  |  |  | *k<sub>cat</sub><sup>S4</sup>* | 3.8 s<sup>-1</sup> | 2.3–6.7 |
|  |  |  | *k<sub>cat</sub><sup>S4</sup>/K<sub>m</sub><sup>S4</sup>* | 12 µM<sup>-1</sup>s<sup>-1</sup> | 7.3–21 |
|  |  |  | *K<sub>m</sub><sup>S5</sup>* | 0.27 µM | 0.056–0.94 |
|  |  |  | *k<sub>cat</sub><sup>S5</sup>* | 3.4 s<sup>-1</sup> | 2.2–7.4 |
|  |  |  | *k<sub>cat</sub><sup>S5</sup>/K<sub>m</sub><sup>S5</sup>* | 13 µM<sup>-1</sup>s<sup>-1</sup> | 7.0–22 |

*k<sub>cat</sub><sup>S2</sup>*/ *K<sub>m</sub><sup>S2</sup>* confidence limits were estimated by KinTek Explorer software. The catalytic efficiencies (*k<sub>cat</sub><sup>SX</sup>* / *K<sub>m</sub><sup>SX</sup>*) where x equals 3-5 were estimated from the corresponding best-fit elementary rate constants.

### Kinetic Comparison Across CRL Family Members Reveals Conserved Catalytic Principles

To ask whether the principles established for CRL2 generalize across the CRL family, substrate priming was analyzed by a neddylated CRL1 assembled from the cullin CUL1, the substrate receptor FBXW7–SKP1, and the RING-between-RING ubiquitin-carrying enzyme ARIH1 (^N8^CRL1^FBXW7^), acting on a phosphorylated CYCLIN E– derived peptide (Supplementary Figure S5a). To restrict the reaction to a single priming event, a ubiquitin variant was employed where all Lys residues had been replaced with Arg (K0 ubiquitin). Reactions were carried out similarly as for the CRL2 system (ARIH1 titration in single-turnover quench-flow assays and steady-state time courses followed by data fitting in KinTek Explorer). While ARIH1 employs the UCE UBE2L3 for thioesterification to ubiquitin (Figure 4a), this step was excluded from the model based on the assumption that it is not rate limiting.

**Figure 4.**
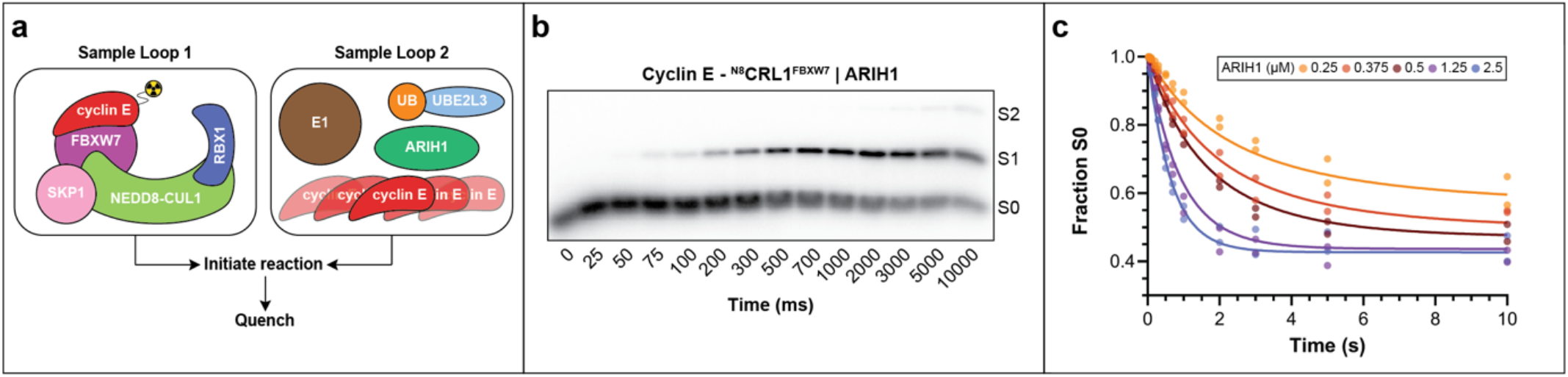
CRL-dependent substrate ubiquitylation modeling is generalizable beyond CRL2. (a) Pre-steady state ubiquitylation reactions with neddylated CRL1^FBXW7^ and radiolabeled Cyclin E peptide (S0) under single-encounter conditions. Ubiquitin-carrying enzymes ARIH1 and UBE2L3, thioesterified to K0 ubiquitin (Lys to Arg mutant to prevent chain formation), were pre-incubated before CRL-S0 mixing. (b) Representative autoradiogram showing S1 product appearance at 2.5 μM ARIH1. (c) S0 loss over time at indicated ARIH1 concentrations, with best-fit lines from KinTek Explorer using the model in Supplementary Figure S5b. Duplicate data points are shown from n=2 technical replicates. Abbreviations: UB, ubiquitin; ^N8^CRL1^FBXW7^, neddylated CUL1-RBX1 in complex with FBXW7-SKP1 substrate receptor.

Global fitting of the data to the model yielded well-constrained rate constants and revealed striking kinetic similarity between CRL1 and CRL2 systems despite their divergent architectures (Figure 4 and Supplementary Figures S5 and S6). Like the CRL2 data, traditional assessment of the kinetic traces showed that the observed exponential rate constants were hyperbolic with respect to ARIH1 levels, suggesting saturation of the CRL at high concentration (Supplemental Figure S5g). Amplitude and ARIH1 also fit well to a hyperbola (Supplemental Figure S5h). The apparent substrate dissociation constants were comparable (*K*_*d,app*_ = 0.5 µM for Hif1α versus 0.3 µM for Cyclin E peptides; Table 2), and the UCE *K*_*m*_ values were statistically identical (1.7 µM for both UBE2R2 and ARIH1). Although the catalytic rates differed modestly (1.0 s^−1^ for UBE2R2 versus 2.6 s^−1^ for ARIH1; Table 1), the resulting catalytic efficiencies differed by less than 3-fold.

### Drug-Induced Neo-substrate Ubiquitylation Displays Atypical Kinetic Properties

CRLs are increasingly exploited therapeutically through targeted protein degradation, in which a small molecule induces proximity between a substrate receptor and a neo-substrate, a protein that the ligase does not normally recognize but is recruited in the presence of the drug (55-60). We reconstituted neo-substrate ubiquitylation with neddylated CUL4–RBX1 bearing the substrate receptor CRBN–DDB1–DDA1 (^N8^CRL4^CRBN^), the validated ubiquitin-carrying enzyme UBE2D3 (61), and pomalidomide, a molecular-glue drug approved for multiple myeloma (62) (Supplementary Figure S7a). The neo-substrate was a SALL4 fragment (residues 378–435) previously demonstrated as a pomalidomide-dependent substrate in tissue culture cells (63). Single-turnover reactions were performed as for the natural-substrate systems: a pre-formed complex of radiolabeled SALL4(378–435), ^N8^CRL4^CRBN^, and pomalidomide was mixed with increasing UBE2D3 levels and excess unlabeled neo-substrate in the rapid quench-flow followed by fitting the data globally in KinTek Explorer.

Single-turnover analysis defined a well-constrained neo-substrate dissociation constant. However, unlike the natural-substrate systems, the UBE2D3 dissociation rate and the ubiquitin-transfer rate only displayed lower limits in FitSpace analysis (Figure 5 and Supplementary Figure S7). Traditional data analysis provided a rationale for this observation: the exponential rate of substrate consumption increased linearly with UBE2D3 concentration rather than saturating (Supplementary Figure S7l), indicating that even the highest enzyme concentrations did not approach saturation of the CRL. Here the ratio *k*_*cat*_/*K*_*m*_ and not *k*_*cat*_ and *K*_*m*_ separately is defined by the experiment.

**Figure 5.**
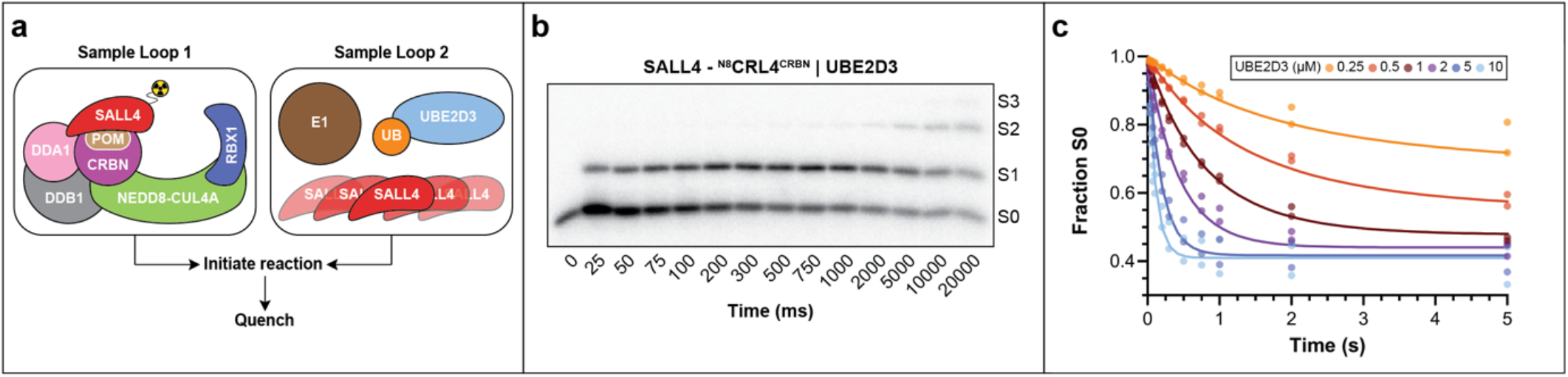
Global data fitting characterizes neo-substrate ubiquitylation in targeted protein degradation. (a) Pre-steady state ubiquitylation reactions with neddylated CRL4^CRBN^ and radiolabeled SALL4(378–435) neo-substrate (S0) under single-encounter conditions. Molecular glue ligand pomalidomide induces CRBN–SALL4 proximity. UBE2D3 was charged with K0 ubiquitin before reaction initiation. (b) Representative autoradiogram showing S1 product appearance at 10 μM UBE2D3. (c) S0 loss over time at indicated UBE2D3 concentrations, with best-fit lines from KinTek Explorer using model in Supplementary Figure S7b. Duplicate data points are shown from n=2 technical replicates. Abbreviations: UB, ubiquitin; ^N8^CRL4^CRBN^, neddylated CUL4A-RBX1 in complex with CRBN-DDB1-DDA1 substrate receptor complex; POM, pomalidomide.

Using a modified modeling approach to extract this ratio (Supplementary Figure S8), a catalytic efficiency of 1.2 µM^−1^s^−1^ was estimated, remarkably close to the values for natural substrates reported here (0.6 –1.6 µM^−1^ s^−1^; Table 2). This near-equivalence of efficiency, however, conceals a fundamentally different partitioning of the underlying constants: the neo-substrate reaction combines a much faster ubiquitin-transfer rate (higher *k*_*cat*_) with a proportionally elevated Michaelis constant (Table 1).

Finally, we note that recent structures of CRLs chemically trapped to capture ubiquitin transfer from UCE to CRL-bound substrates have shown potential interfaces between the UCE, the substrate and the CRL substrate receptor (17,18,31,32,34). To test whether reaction components such as the UCE affected binding, the kinetics of neo-substrate association and dissociation were measured by stopped flow fluorescence anisotropy experiments, using AlexaFluor568-labeled SALL4(378–435) and the isolated substrate-receptor complex (Figure 6a and Supplementary Figure S9). Global fitting gave an association rate of 2.1 µM^−1^ s^−1^ and a dissociation rate of 0.290 s^−1^, corresponding to a *K*_*d*_ of 0.14 µM (Figure 6b–f), only about 2-fold tighter than the value estimated from the full kinetic reaction (Table 2). This modest difference indicates that the neddylated cullin and the ubiquitin-carrying enzyme may slightly weaken neo-substrate affinity for the substrate-receptor complex, but do not substantially perturb binding.

**Figure 6.**
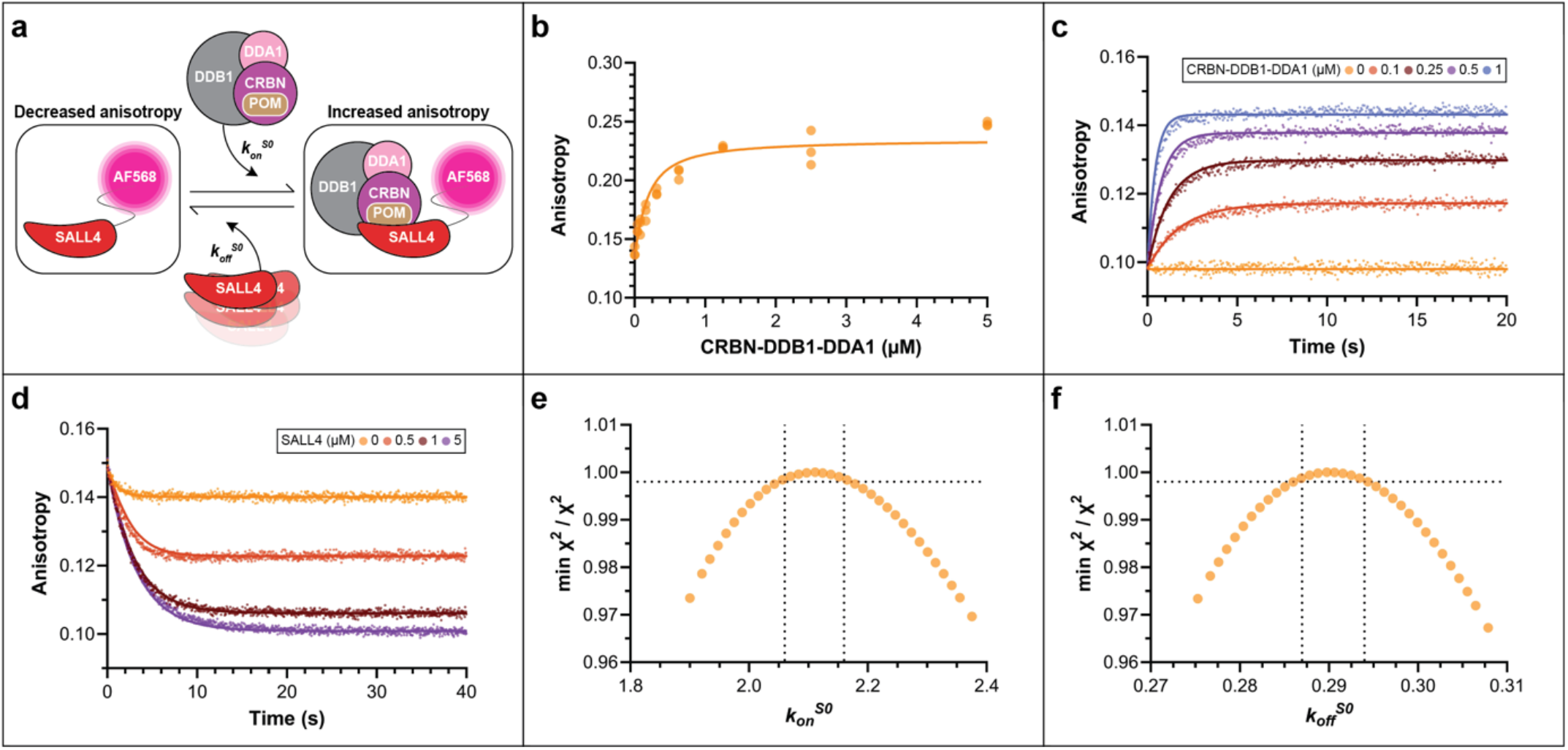
Global fitting reveals distinct binding kinetics between neo-substrate and CRBN from fully reconstituted reactions. (a) Fluorescence polarization-based binding assay between SALL4(378–435) neo-substrate and CRBN-DDB1-DDA1 substrate receptor complex enables both equilibrium and kinetic binding studies. (b) Equilibrium binding curve for AlexaFluor568-labeled SALL4(378–435) in the presence of pomalidomide and increasing substrate receptor concentrations, fit using the model in Supplementary Figure S9b via KinTek Explorer. (c) Pre-steady state association kinetics: fluorescently-labeled SALL4(378–435) mixed with increasing CRBN-DDB1-DDA1 concentrations in the presence of pomalidomide, analyzed by global data fitting. (d) Dissociation kinetics: pre-formed AlexaFluor568-labeled SALL4(378–435)–substrate receptor complex diluted with excess unlabeled SALL4(378–435). (e–f) Confidence contour plots for *k*_*on*_^*S0*^ (e) and *k*_*off*_^*S0*^ (f) rate constants showing 95% confidence limits. Abbreviations: POM, pomalidomide; AF568, AlexaFluor568 fluorescent dye. Equilibrium experiments: n=3 technical replicates; pre-steady state binding kinetics: n=8 technical replicates.

## Discussion

By fitting families of pre-steady-state and steady-state measurements globally to explicit kinetic models, and testing each model by confidence-contour analysis, we have resolved several features of CRL-dependent ubiquitylation that conventional steady-state treatments leave hidden. Each pre-steady-state trace harbors two independent pieces of information and separating them is central to the analysis. The observed rate is an eigenvalue, defined as a combination of the underlying microscopic rate constants, whereas the amplitude reports the position of an equilibrium that is established before the rate-limiting step. In the priming experiments (Figure 1), the dependence of the amplitude on enzyme concentration reports on the partitioning of substrate between free and CRL-bound states, explaining why the amplitude data constrain the substrate dissociation constant while the eigenvalues report on chemistry. This is evident through simulation in KinTek Explorer: exclusion of the CRL-S0 binding step from the priming model results in each trace corresponding to increasing UBE2R2 concentration eventually achieving the same endpoint.

Below we consider what the method makes measurable, what the conserved efficiencies imply about CRL catalysis, and where the analysis remains incomplete.

### What Global Data Fitting with KinTek Explorer Offers Over Previous Models for Protein Ubiquitylation

A recurring difficulty with conventional, equation-based kinetic analysis is that it can return apparently precise rate constants that the data do not constrain. Standard errors from nonlinear regression are often overoptimistic and give little warning when a parameter is bounded only on one side. Fitting the complete time courses by numerical integration and then mapping each parameter with confidence-contour (FitSpace) analysis provides an explicit statement of which constants are well constrained and their confidence intervals defined by upper or lower limits (52).

Neo-substrate ubiquitylation by ^N8^CRL4^CRBN^ illustrates the point concretely. Conventional analysis by nonlinear curve fitting to a double-exponential solution previously used to model ubiquitylation (46) returned a rate of ubiquitin transfer (*k*_*obs*_) with deceptively tight error bounds (Supplementary Figure S10), yet confidence-contour analysis showed that this rate, as well as the UBE2D3 dissociation rate, were constrained only by lower limits (Supplementary Figure S7j,k). This was also apparent when examining the exponential rate of substrate consumption which increases linearly with UBE2D3 concentration rather than trending toward saturation (Supplementary Figure S7l). Thus, only the ratio *k*_*cat*_/*K*_*m*_, not the two constants individually, is defined by the experiment.

A second advantage of global data fitting and our model is the ability to estimate substrate affinity (*K*_*d,app*_) for the assembled CRL from kinetic data alone, without appending a bulky fluorophore that may itself perturb binding. Across three systems, global fitting placed the apparent substrate dissociation constants within a narrow, sub-micromolar window (0.24 µM for the SALL4 neo-substrate with ^N8^CRL4^CRBN^, 0.49 µM for Hif1α peptide with ^N8^CRL2^VHL^, and 0.29 µM for phosphorylated Cyclin E peptide with ^N8^CRL1^FBXW7^). Because these constants are extracted from the fully reconstituted reaction rather than from isolated components, they describe substrate binding in the catalytic environment. Moreover, the results show that the dissociation rates are slow enough relative to the estimated ubiquitin transfer rates for the complex to persist through priming and successive chain-extension events.

Whether the other reaction components shape affinity between substrate and CRL can be tested directly. For the neo-substrate SALL4 and the isolated CRL substrate receptor CRBN, a fluorescence binding assay yielded a *K*_*d*_ of 0.14 µM, within 2-fold of the estimate from the fully assembled complex, indicating that the neddylated cullin and UBE2D3 do not substantially alter neo-substrate binding in this system. It is not expected that this equivalence will hold universally since UCEs that build poly-ubiquitin chains, including UBE2R-family members and UBE2G1, carry a binding surface for substrate-linked ubiquitin and may therefore tighten substrate affinity for the CRL–UCE∼ubiquitin complex as a chain grows (32,61).

Finally, we note that modeling consecutive ubiquitin transfer events by conventional methods is notoriously difficult, owing to the necessity of extracting individual rates for each transfer cycle. This requires fitting each time course for each product species to a sum of exponentials, resulting in inflation of fitting errors for each additional ubiquitin transfer. Global simulation-based fitting of all species simultaneously avoids this problem by constraining every intermediate with the same set of microscopic rate constants.

### Catalytic Efficiencies Reveal Conserved CRL Strategies Despite Architectural Diversity

Resolving *k*_*cat*_/*K*_*m*_ for priming across three architecturally distinct CRLs allows for comparison of their catalytic efficiencies directly, whereas prior literature relied on *K*_*m*_ estimates derived from steady-state experiments that may inaccurately suggest saturation of the CRL owing to a lack of time resolution during data collection. Despite major differences in substrate receptor (F-box for CUL1, VHL for CUL2, CRBN for CUL4), in substrate, and in the partner UCE, the priming efficiencies span only 3-fold, from 0.6 µM^−1^ s^−1^ for UBE2R2 with ^N8^CRL2^VHL^ to 1.6 µM^−1^ s^−1^ for ARIH1 with ^N8^CRL1^FBXW7^.

This convergence is notable: ARIH1 is an RBR-type E3 that primes through a hybrid RING/HECT mechanism (39,64) and is distinct in size and fold in comparison with the E2s UBE2R2 and UBE2D3. Even more remarkable is that a pomalidomide-recruited SALL4 fragment achieves a priming efficiency of 1.2 µM^−1^ s^−1^, squarely within the natural-substrate range despite no component of this reaction being under selective pressure to ubiquitylate this target. As such, it seems that CRL-UCE catalytic efficiency is an intrinsic property of the shared cullin–RBX catalytic geometry (31) and that human CRLs function with multiple distinct UCEs, allowing for rapid ubiquitylation despite substrate heterogeneity in shape, size and Lys presentation.

### Advancing the Kinetic Framework

While our analysis represents a substantial advance in quantifying CRL-dependent ubiquitylation, the mechanistic framework remains open to refinement. Model complexity must be commensurate with the information content of the data, requiring compromises during modeling that included assuming diffusion-driven kinetics for several steps and the linking of rate constants that may affect modeling in distinct ways. For instance, regarding the binding of UCE to the CRL complex, locking the rate of association has the consequence that the magnitude of the rate of UCE dissociation is unknown, and instead their ratio, the *K*_*m*_ of UCE for the CRL, is estimated (52). Similarly, our models require that the UCE is in rapid equilibrium with the CRL, which has been directly validated for UBE2R2 (51) but does not inform on exactly how fast UCE binds. Furthermore, rapid equilibrium has not been tested for ARIH1 and its partner UCE UBE2L3 or UBE2D3 (see below). Finally, there are cases where UCE saturation of the CRL was not achievable, resulting in the model providing *k*_*cat*_/*K*_*m*_, such as in the case of UBE2D3 and ^N8^CRL4^CRBN^, with only lower limits on *k*_*cat*_ and *K*_*m*_ individually rather than discrete values for these parameters.

Future stopped-flow kinetic studies could directly measure UCE binding to the CRL and dissociation rates with millisecond temporal resolution, testing model predictions regarding UCE dynamics. Single-molecule fluorescence approaches could reveal whether individual CRL-UCE pairs undergo conformational changes during catalysis or whether the ensemble kinetics we observe masks heterogeneity in catalytic mechanisms. Comprehensive analysis of the full CRL family (CUL3, CUL5, CUL7, CUL9) would establish whether the catalytic efficiency conservation we observe extends universally or whether subfamily-specific mechanisms exist. Finally, we note that while the quantitative insights here facilitate comparative analysis of physiological ubiquitylation mechanisms, they equally apply to design of proximity-inducing drugs for targeted protein degradation therapeutics where optimization of CRL-substrate-drug interactions could accelerate ubiquitin transfer that may enhance degradation efficiency in therapeutic applications.

## Acknowledgements

We thank Kleiger lab members Tanjina Tarannum, Pooja Verma, Chutima Rattanasopa and Isabella Romero Yusti for helpful discussions and the sharing of reagents and laboratory protocols and Casey Hall and the UNLV Genomics Core for DNA sequencing. This research was supported by NIH R01 GM141409, NCI R01 CA279255, and NIH R35 GM163591 (N.P., J.L., and G.K.). B.A.S. was supported by the Max Planck Society. The content is solely the responsibility of the authors and does not necessarily represent the official views of the National Institutes of Health. This preprint was typeset with the bioRxiv word template by @Chrelli: www.github.com/chrelli/bioRxiv-word-template

## Author contributions

N.P., J.L., T.L.D. and G.K. conceived the project. N.P., J.L. and D.C.S. generated and purified proteins; D.C.S. was supervised by B.A.S. N.P. and J.L. performed the enzymology and were supervised by G.K. T.L.D. and G.K. designed the kinetic models and performed data fitting and confidence contour analysis in KinTek Explorer. N.P. prepared all figures with input from G.K. G.K. and T.L.D. prepared the manuscript with input from all authors. G.K. supervised the project.

## Competing interest statement

B.A.S. is a member of the scientific advisory boards of Proxygen and Lyterian and has financial interest in Nextech Invest. The other authors declare no competing interests.

**Supplementary Figure S1.**
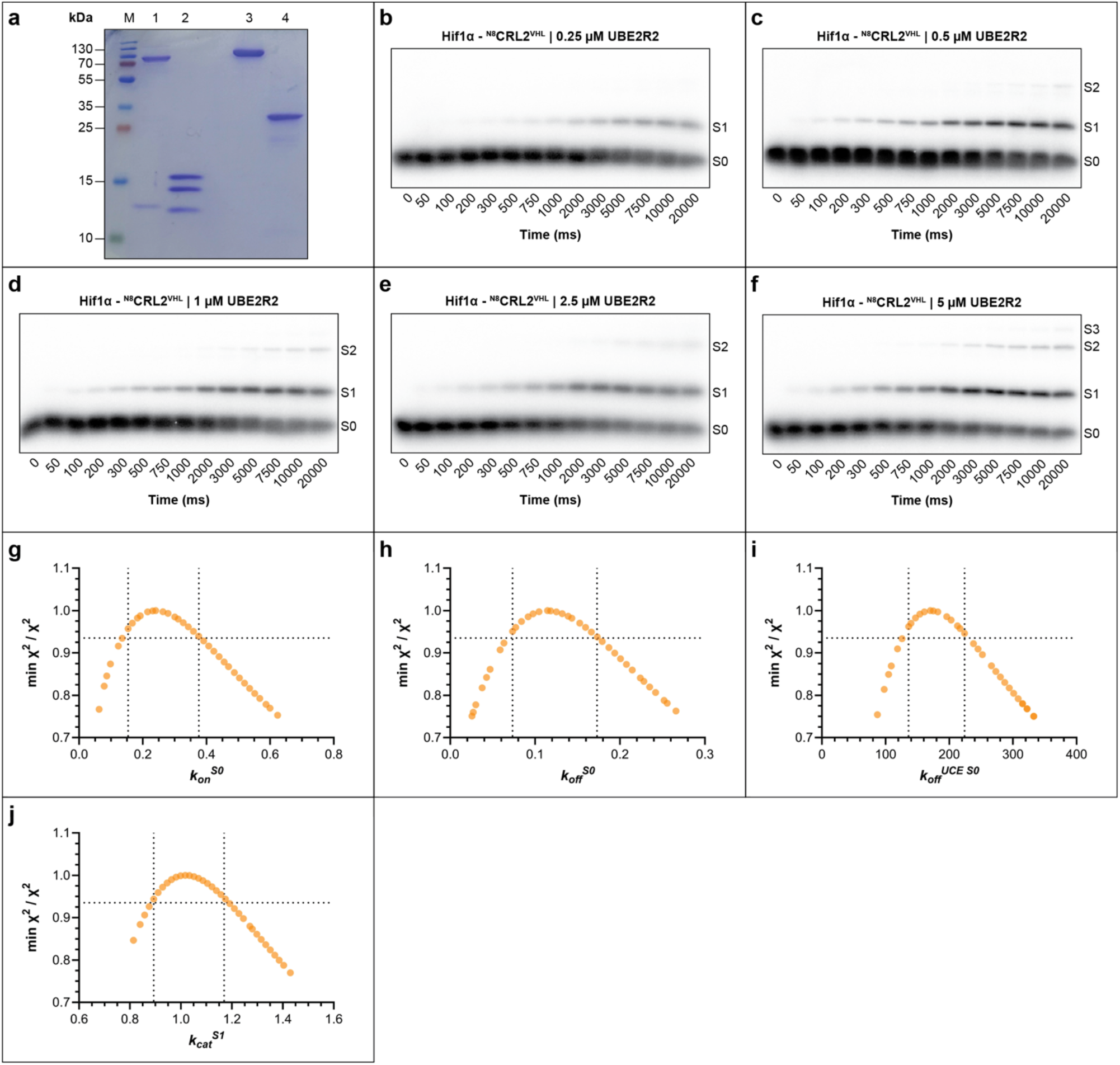
*In vitro* reconstitution and pre-steady state kinetics of CRL2-UBE2R2 catalyzed substrate priming. (a) SDS-PAGE showing the purity levels of various enzyme and protein components. Lane M shows molecular weight markers, lane 1 neddylated CUL2-RBX1, lane 2 VHL in complex with ELONGIN B and ELONGIN C, lane 3 E1, and lane 4 UBE2R2. (b) Autoradiogram showing primary conversion of S0 substrate to S1 product at 0.25 μM UBE2R2. (c) Same as (b), except with 0.5 μM UBE2R2. (d) same as (b), except with 1 μM UBE2R2. (e) Same as (b), except with 2.5 μM UBE2R2. (f) Same as (b), except with 5 μM UBE2R2. (g–j) Confidence contour plots showing (χ^2^ minimum/χ^2^) values for the indicated rate constants across a range of values, where 1 represents the best fit. Horizontal and vertical dotted lines indicate 95% confidence limits. n=2 technical replicates.

**Supplementary Figure S2.**
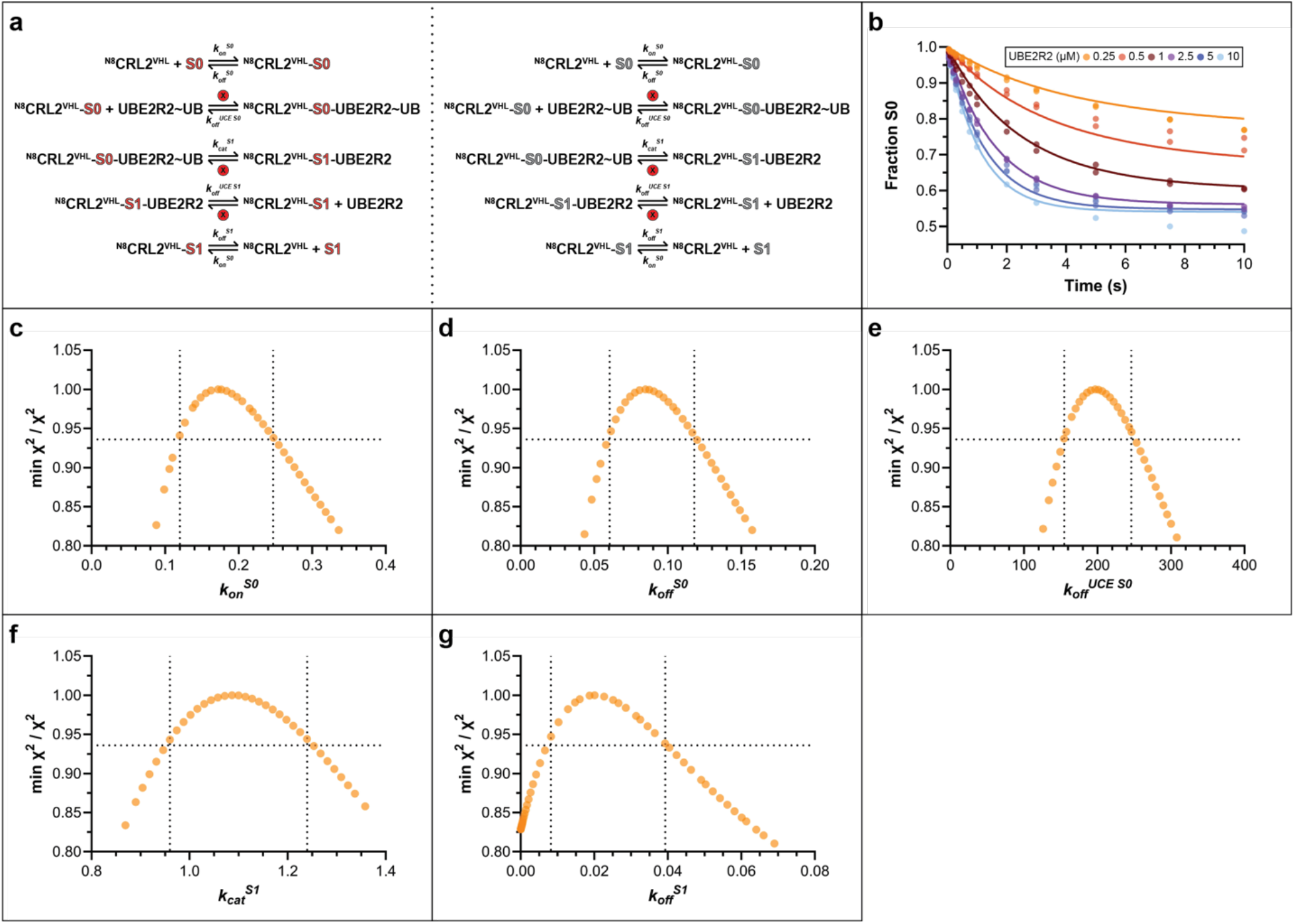
Expanded kinetic model incorporating product association and dissociation rates from ^N8^CRL2^VHL^. (a) Kinetic model similar to Figure 1e that also includes a rate constant for S1 product dissociation (*k*^*offS1*^). Circled ‘x’ indicates rate constants with locked values. Steps with unlabeled excess S0 (grey) promote single-encounter conditions between radiolabeled substrate and the CRL. (b) S0 disappearance as a function of time and UBE2R2 concentration, with best-fit lines from KinTek Explorer based on the model in (a). (c-g) Confidence contour plots showing (χ^2^ minimum/χ^2^) values for the indicated rate constants across a range of values, where 1 represents the best fit. Horizontal and vertical dotted lines indicate 95% confidence limits.

**Supplementary Figure S3.**
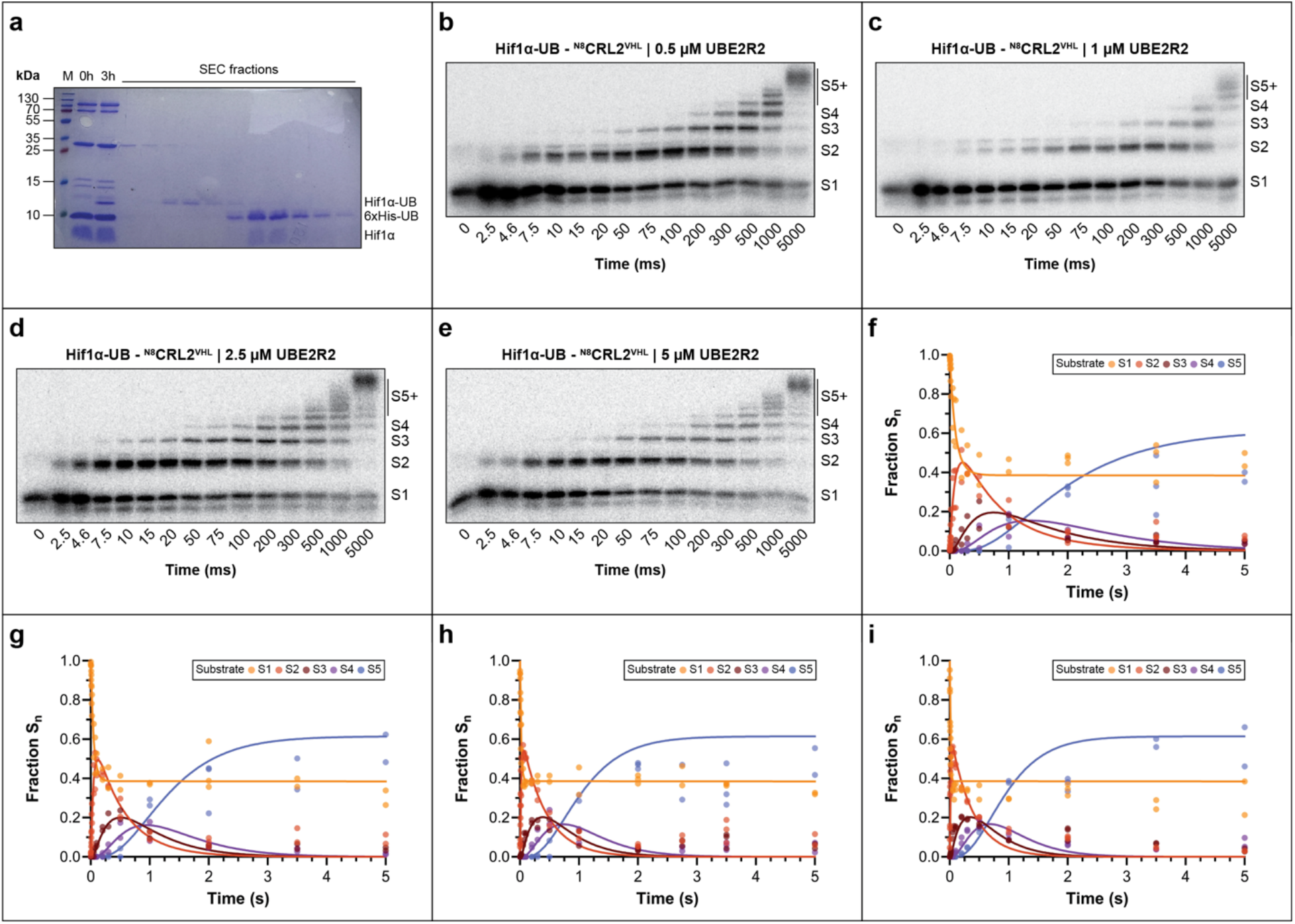
Pre-steady state kinetics and global data fitting of consecutive ubiquitin transfer events and poly-ubiquitin chain formation. (a) SDS-PAGE showing purification of mono-ubiquitylated Hif1α peptide (S1). Lane M shows molecular weight markers, lanes 1 and 2 show reaction components and S1 product formation after 3 hours, respectively. Additional lanes show fractions from size exclusion chromatography. Note that highly pure S1 eluted prior to unconjugated ubiquitin and Hif1α peptide (S0). (b) Representative autoradiogram at 0.5 μM UBE2R2 showing S1 conversion to poly-ubiquitylated products. All non-overlapping product bands were quantified. (c) Same as (b), except with 1 μM UBE2R2. (d) Same as (b), except with 2.5 μM UBE2R2. (e) Same as (b), except with 5 μM UBE2R2. (f) S1 loss and appearance of products S2–S5 with best-fit lines from KinTek Explorer and 0.5 μM UBE2R2. (g) Same as (f), except with 1 μM UBE2R2. (h) Same as (f), except with 2.5 μM UBE2R2. (i) Same as (f), except with 5 μM UBE2R2. n=2 technical replicates.

**Supplementary Figure S4.**
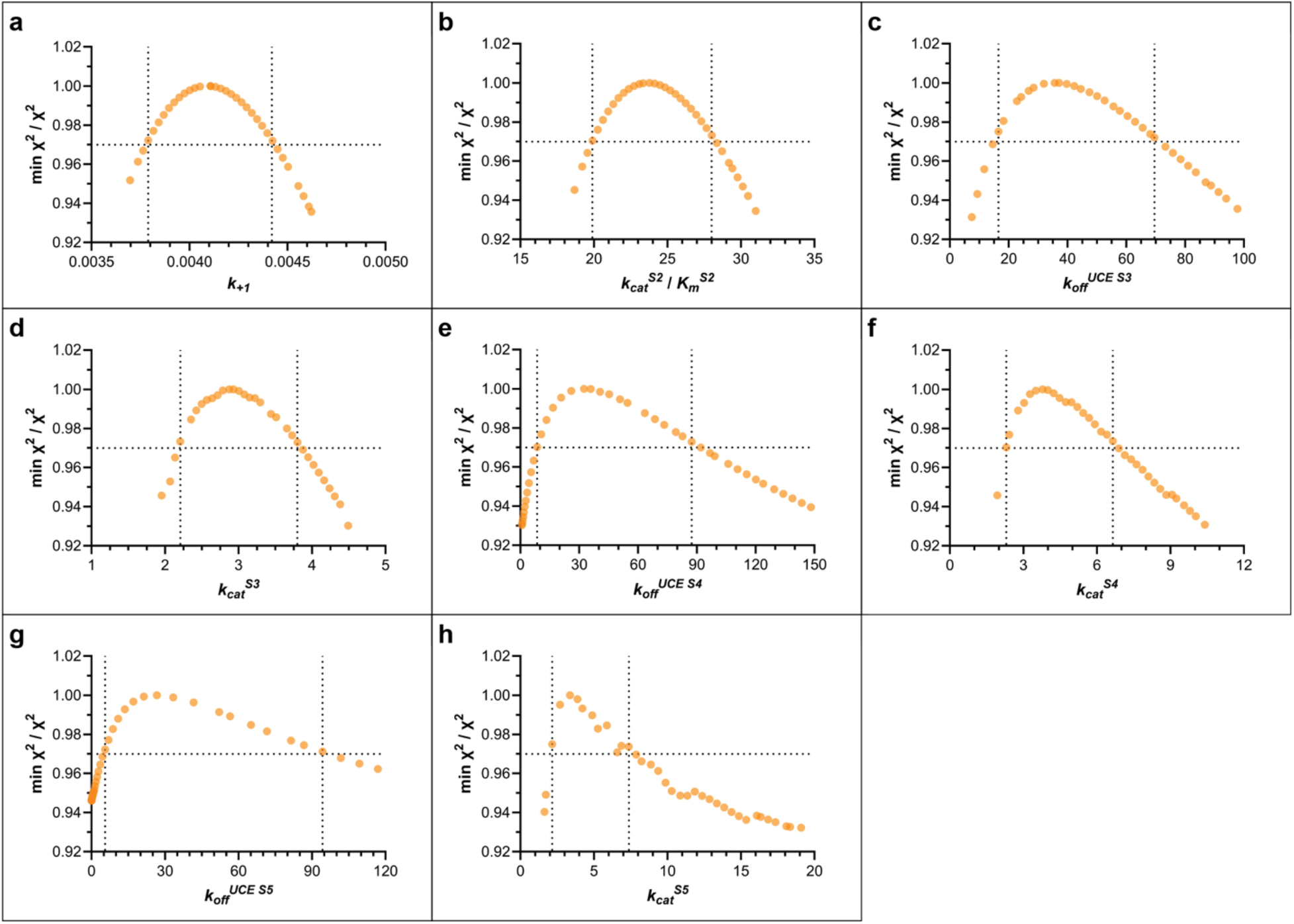
Confidence-contour analysis through FitSpace in KinTek Explorer shows that all floated rate constants are constrained by the data for poly-ubiquitin chain formation. (a-h) Confidence contour plots showing (χ^2^ minimum/χ^2^) values for the indicated rate constants across a range of values, where 1 represents the best fit. Horizontal and vertical dotted lines indicate 95% confidence limits.

**Supplementary Figure S5.**
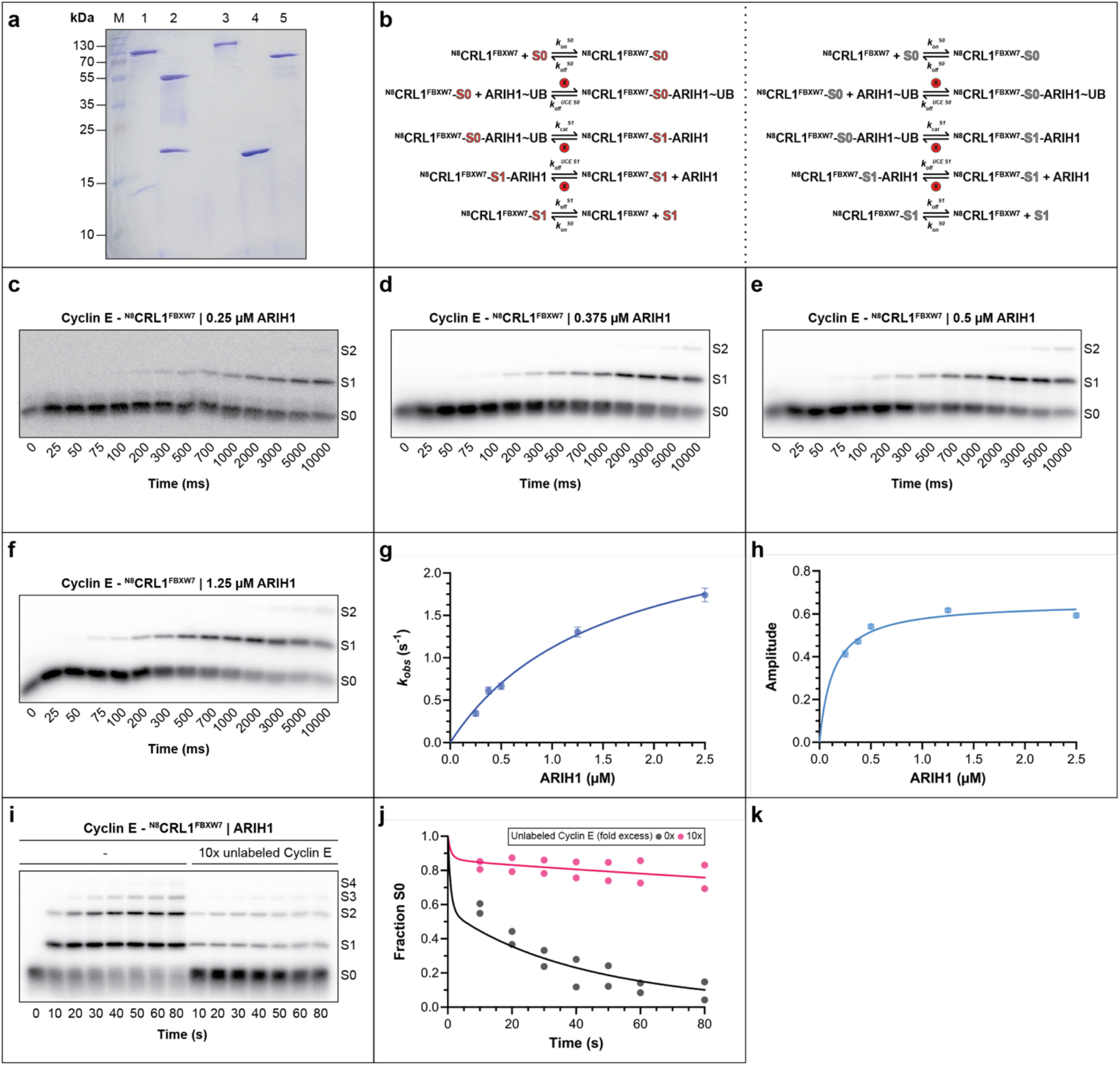
*In vitro* reconstitution and pre-steady state kinetics of CRL1-ARIH1 catalyzed substrate priming enable model comparison across multiple CRL sub-families. (a) SDS-PAGE showing the purity levels of various enzyme and protein components. Lane M shows molecular weight markers, lane 1 neddylated CUL1-RBX1, lane 2 truncated FBXW7 in complex with SKP1, lane 3 E1, lane 4 UBE2L3 and lane 5 ARIH1. (b) Kinetic model similar to Supplementary Figure S2a, except for ARIH1-catalyzed substrate priming with ^N8^CRL1^FBXW7^ and Cyclin E peptide substrate. Circled ‘x’ indicates rate constants with locked values. Steps with unlabeled excess S0 (grey) promote single-encounter conditions between radiolabeled S0 and CRL. (c) Representative autoradiogram showing S1 product appearance at 0.25 μM ARIH1. (d) Same as (c), except with 0.375 μM ARIH1. (e) Same as (c), except with 0.5 μM ARIH1. (f) same as (c), except with 1.25 μM ARIH1. (g) Graph showing the exponential observed rate constant (*k*_*obs*_) for S0 disappearance as a function of ARIH1 concentration. (h) Graph showing the amplitude of S0 depletion at equilibrium as a function of ARIH1 concentration. (i) Autoradiogram for single-turnover (left half) and multi-turnover (right half) reactions. (j) Substrate (S0) depletion kinetics and the fit of the data to the model for single-turnover (grey) and multi-turnover (magenta). n=2 technical replicates.

**Supplementary Figure S6.**
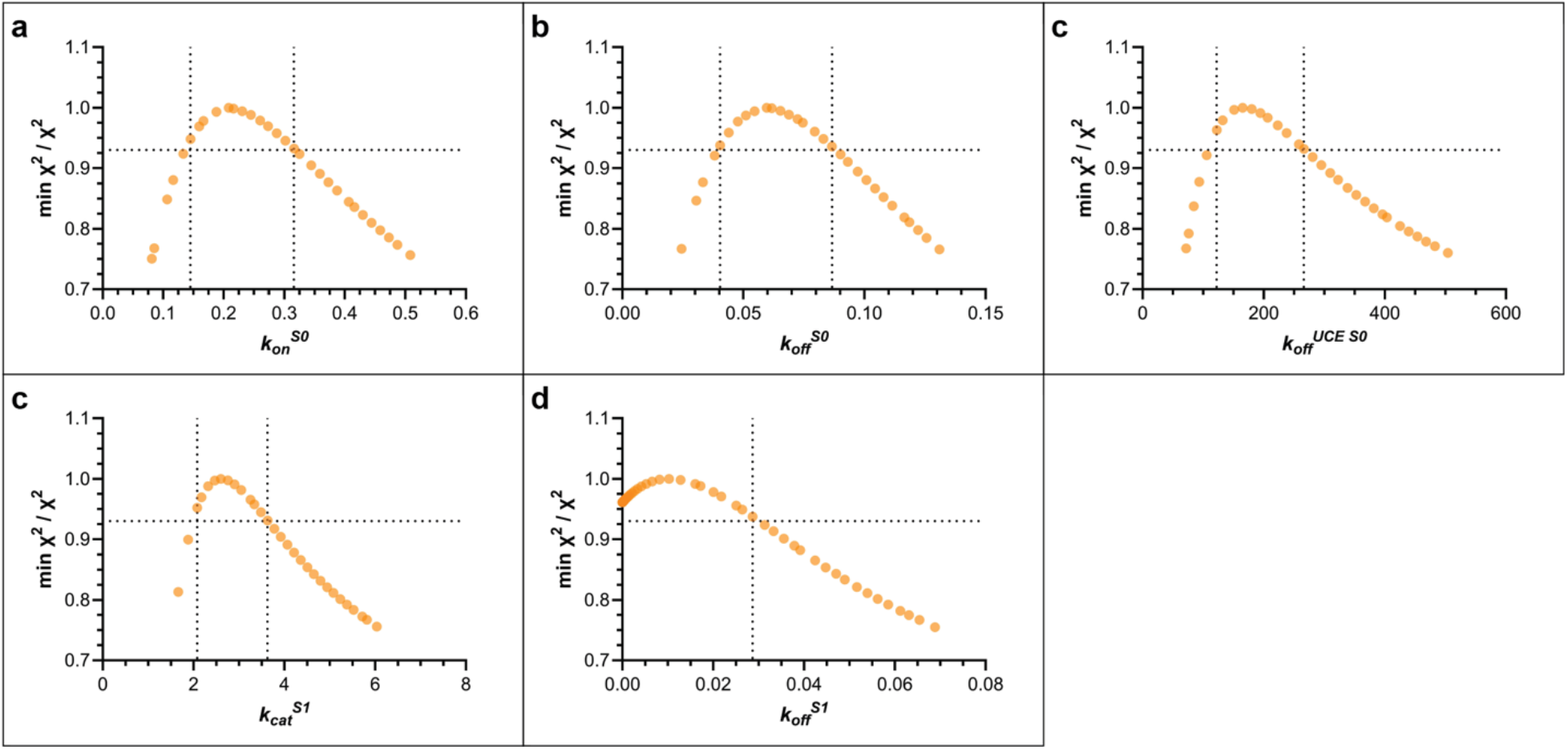
Confidence-contour analysis for CRL1-ARIH1 catalyzed substrate priming show well-constrained rate constants. (a-e) Confidence contour plots showing (χ^2^ minimum/χ^2^) values for the indicated rate constants across a range of values, where 1 represents the best fit. Horizontal and vertical dotted lines indicate 95% confidence limits.

**Supplementary Figure S7.**
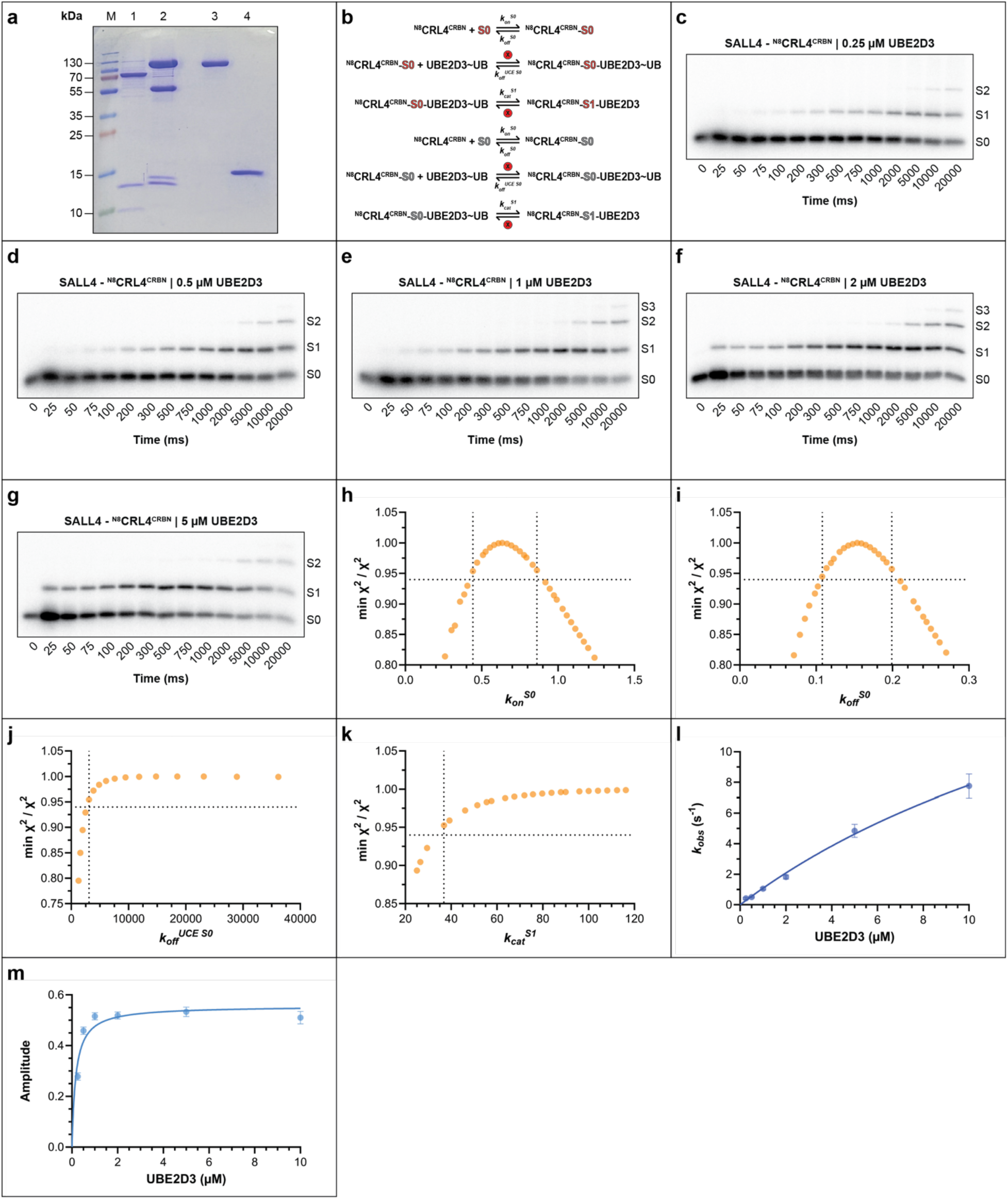
*In vitro* reconstitution and pre-steady state kinetics of a neo-substrate of targeted protein degradation shows distinct kinetics compared with natural substrates. (a) SDS-PAGE showing the purity levels of various enzyme and protein components. Lane M shows molecular weight markers, lane 1 neddylated CUL4A-RBX1, lane 2 CRBN with DDB1 and DDA1, lane 3 E1, and lane 4 UBE2D3. (b) Kinetic model similar to Figure 1e except for UBE2D3-catalyzed priming with ^N8^CRL4^CRBN^ and SALL4(378–435) neo-substrate (S0). Circled ‘x’ indicates rate constants with locked values. Steps with unlabeled excess S0 (grey) promote single-encounter conditions between radiolabeled substrate and CRL. (c) Representative autoradiogram showing S1 product appearance at 0.25 μM UBE2D3. (d) Same as (c), except with 0.5 μM UBE2D3. (e) Same as (c), except with 1 μM UBE2D3. (f) same as (c), except with 2 μM UBE2D3. (g) same as (c), except with 5 μM UBE2D3. (h-k) Confidence contour plots showing (χ^2^ minimum/χ^2^) values for the indicated rate constants across a range of values, where 1 represents the best fit. Horizontal and vertical dotted lines indicate 95% confidence limits. (l) Graph showing the exponential observed rate constant (*k*_*obs*_) for S0 disappearance as a function of UBE2D3 concentration. (m) Same as (l), except for the amplitude of fraction S0. n=2 technical replicates.

**Supplementary Figure S8.**
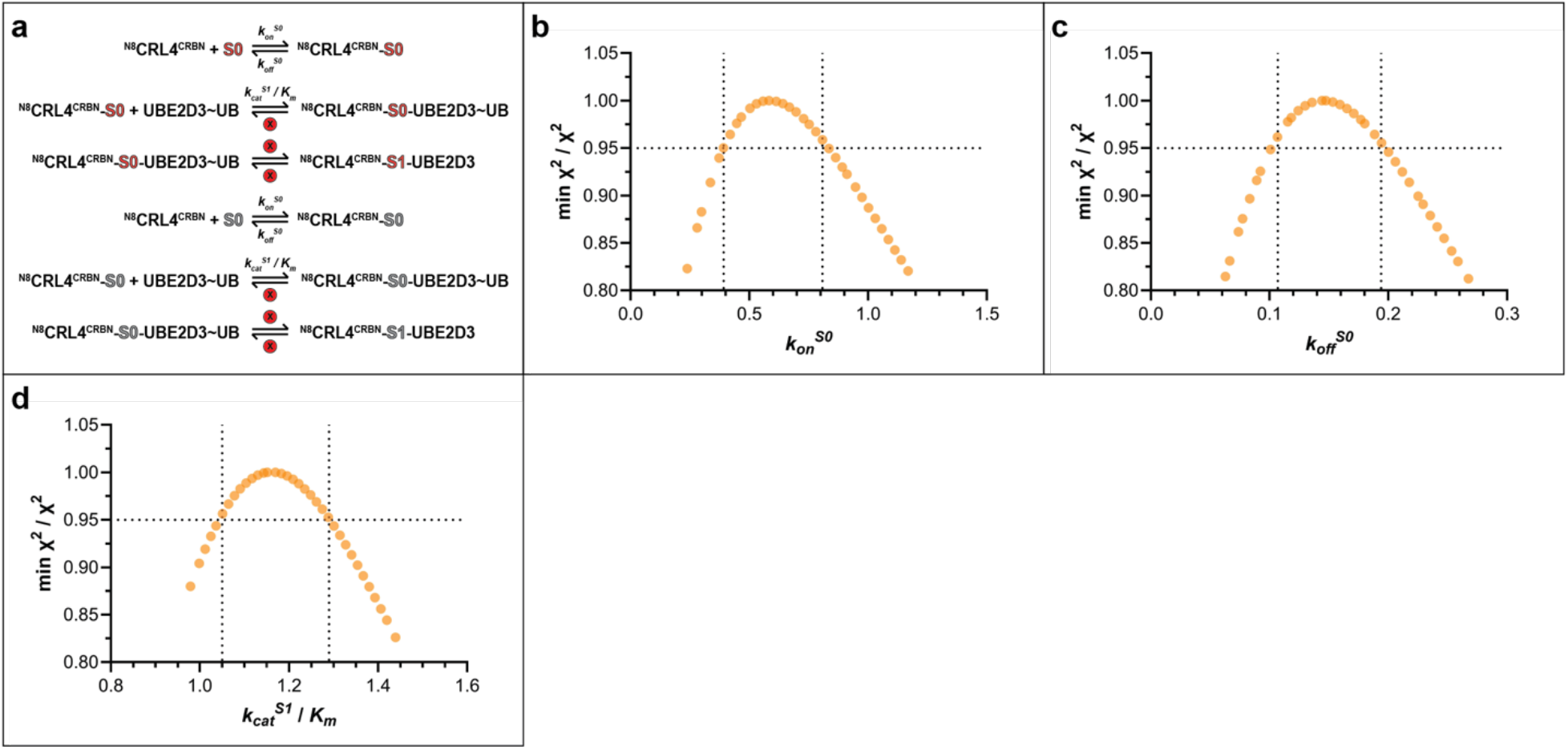
Modified kinetic model allows for estimating the catalytic efficiency of UBE2D3-dependent neo-substrate ubiquitylation. (a) Kinetic model where locking the rate constant for ubiquitin transfer (*k*_*cat*_^*S1/*^) to a high value and the constant for dissociation of UBE2D3∼ubiquitin (*k*_*off*_^*UCE S0*^) to 0 approximates the rate of UBE2D3∼ubiquitin association with CRL as *k*_*cat*_^*S1*^/*K*_*m*_. Circled ‘x’ indicates rate constants with locked values. Steps with unlabeled excess S0 (grey) promote single-encounter conditions between radiolabeled S0 and CRL. (b-d) Confidence contour plots showing (χ^2^ minimum/χ^2^) values for the indicated rate constants across a range of values, where 1 represents the best fit. Horizontal and vertical dotted lines indicate 95% confidence limits.

**Supplementary Figure S9.**
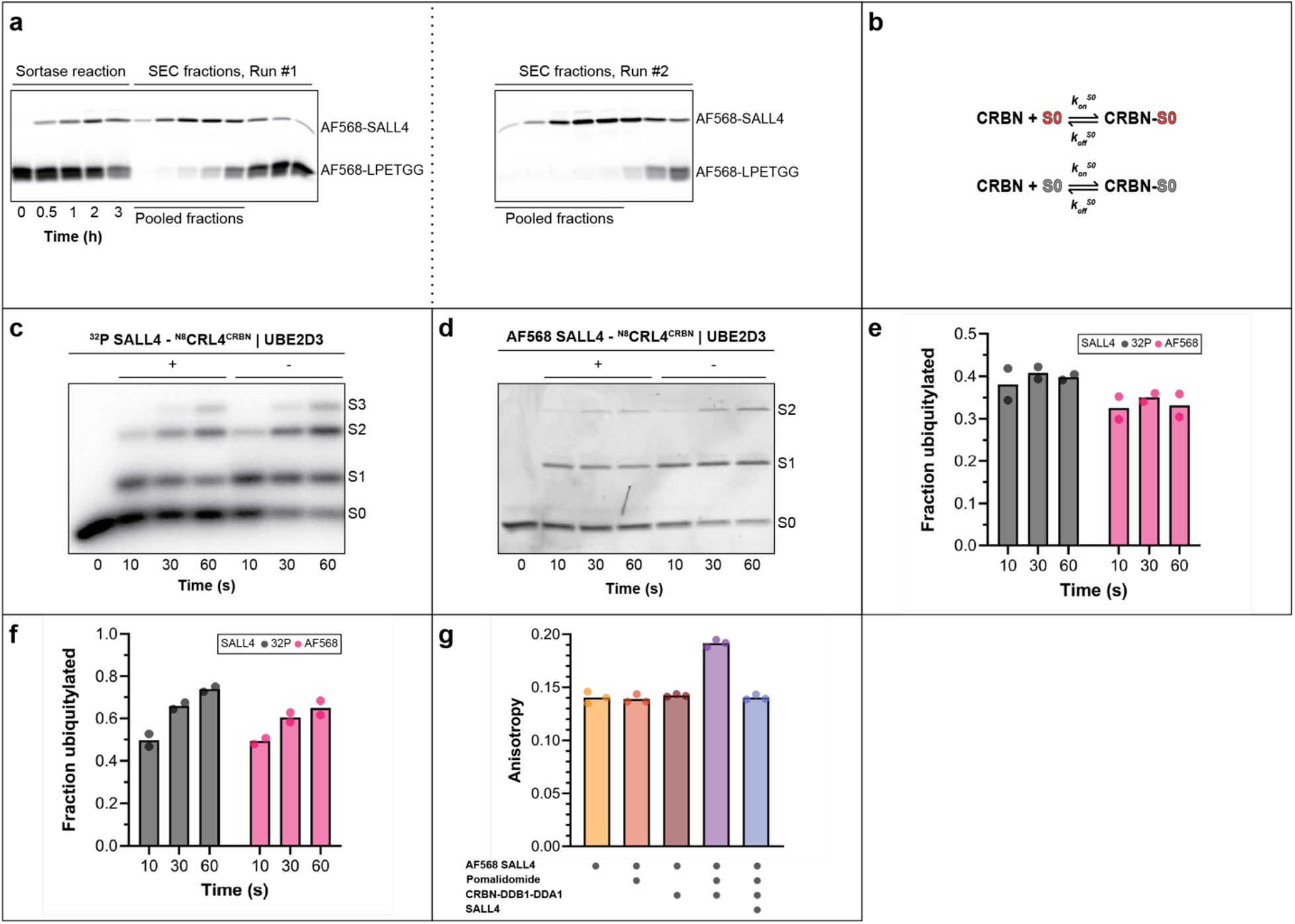
A fluorescence polarization binding assay to estimate the kinetics of neo-substrate binding to CRBN-DDB1-DDA1 substrate receptor complex. (a) Fluorescence scan showing sortase-mediated attachment of an AlexaFluor568-conjugated peptide to SALL4(378–435). The sortase reaction was incubated for 3 hours followed by two rounds of size exclusion chromatography. The right-most scan shows pooled fractions and the purity of labeled SALL4 product that was used for binding experiments. (b) Kinetic model showing rate constants for fluorescently labeled S0 (red) binding to the CRBN-DDB1-DDA1 substrate receptor complex. Unlabeled excess S0 (grey) was included in stopped-flow experiments to estimate the rate of dissociation of labeled substrate from the substrate receptor complex. (c) Ubiquitylation reaction time course showing product formation in the absence (-) or presence (+) of single-encounter kinetics between labeled substrate and CRL. (d) same as (c), except with AlexaFluor568-labeled S0 neo-substrate. (e) Graph comparing product formation for radiolabeled and fluorescently labeled S0 in the presence of single-encounter kinetics. (f) Same as (e) except without single-encounter. (g) Fluorescence polarization signal for binding reactions containing AlexaFluor568-labeled S0, pomalidomide, substrate receptor complex, and competitor unlabeled SALL4. Notice that anisotropy increases when labeled SALL4 is in the presence of substrate receptor and pomalidomide and that this signal can be chased by the addition of competitor substrate. n=3 technical replicates.

**Supplementary Figure S10.**
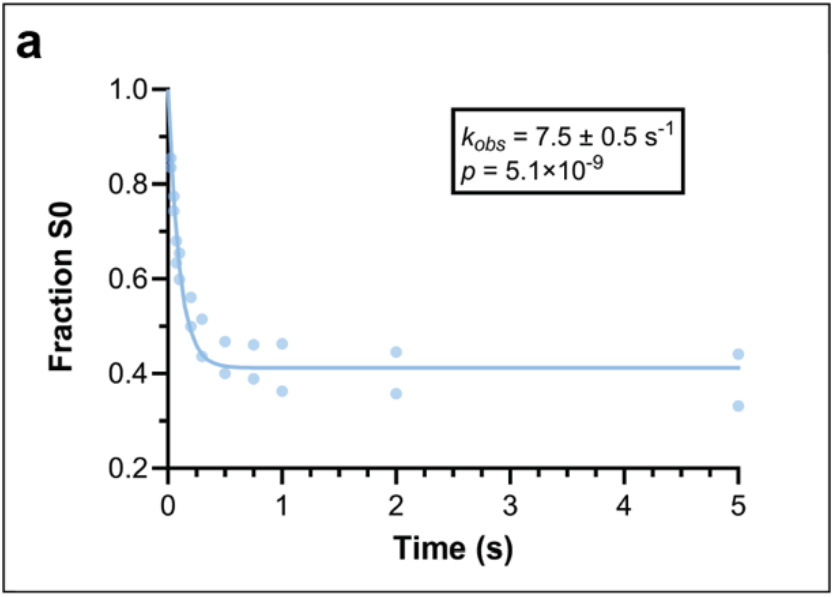
Nonlinear curve fitting to a closed form solution estimates a well-constrained rate of ubiquitin transfer for UBE2D3 and ^N8^CRL4^CRBN^. Results from the 10 μM UBE2D3 dataset showing SALL4(378– 435) neo-substrate (S0) depletion over time (Figure 5b) were fit to a previously described model using Mathematica (46). Notice that the standard error of estimate was less than 7% of the estimated value, whereas global data fitting in KinTek Explorer demonstrated that the rate of ubiquitin transfer was only bounded by a lower limit (Supplementary Figure S7k).

